# Methanogenic archaea bolster mucosal homeostasis and protect from colitis

**DOI:** 10.64898/2026.01.19.700430

**Authors:** Emily E. Byrd, Benjamin W. Harris, Hongpan Zhang, W. Elise Warren, Sree H. Kolli, Audrey M. Putelo, Mitchell T. McGinty, Michael D. Solga, Melanie R. Rutkowksi, Melissa M. Kendall

**Affiliations:** Department of Microbiology, Immunology, and Cancer Biology, University of Virginia School of Medicine, Charlottesville, VA, USA; University of Virginia School of Medicine; Department of Genomic Sciences, University of Virginia School of Medicine, Charlottesville, VA, USA; Flow Cytometry Core Facility, University of Virginia School of Medicine, Charlottesville, VA, USA

## Abstract

The intestinal microbiota profoundly shapes host homeostasis and disease, yet the role of Archaea, which comprise ∼10% of the anaerobic gut microbiota, is poorly understood. Here, we find that colonization of germ-free or conventional mice with the human methanogens *Methanobrevibacter smithii* (MSm) or *Methanosphaera stadtmanae* (MSt) enhances epithelial regeneration, mucus production, and regulatory T cell accumulation. These methanogens protect against chemically induced colitis, demonstrating a direct and beneficial role for Archaea in maintaining colonic health.

---

The gastrointestinal tract harbors trillions of microbes collectively referred to as the microbiota, which promote human health by aiding in intestinal development and nutrient metabolism and defending against pathogens ^1^. However, increasing evidence indicates that the microbiota exacerbates gastrointestinal, cardiovascular, and neurological diseases ^2-4^. The microbiota comprises bacteria, fungi, viruses, and archaea. Methanogenic archaea account for ∼10 % of the anaerobic gut microbiota, and in humans are dominated by five orders ^5^. Of these, MSm and MSt are the most prevalent, with MSm detected in nearly 100% of individuals, and MSm ∼30–90 % respectively ^5,6^. Although correlative and contradictory studies have linked MSm and MSt to inflammatory bowel disease and colorectal cancer ^5,7^, whether these methanogens exert direct effects on colonic physiology and inflammation has remained unanswered.

This knowledge gap is due in large part because culturing methanogens remains technically challenging. Methanogens are strict anaerobes and require oxygen-free conditions for growth. To address this, we cultivated MSm and MSt under strict anaerobic (Hungate-style) conditions ^8,9^, enumerated cells via by DNA staining and F_420_ fluorescence using flow cytometry ^10^, and engrafted either methanogen into germ-free (GF) or antibiotic-treated, specific-pathogen-free (SPF) C57BL/6 mice via two oral gavages (day 0, day 4) (Fig. 1a). Fecal qPCR and culture confirmed that MSm and MSt efficiently colonized the colons of GF and SPF mice (Extended Data Fig. 1a-b), preferentially localizing to the mucus layer in GF mice (Extended Data Fig. 1c-d). As a control, the anaerobic bacterium *Desulfovibrio piger* (DPi), which occupies a similar niche ^11^, was also introduced into GF and SPF mice (Fig. 1a, Extended Data Fig. 1e). At 9 days post-engraftment (dpe), methane levels in cultured fecal slurries were elevated in MSm- and MSt-colonized mice (Fig. 1b), indicating viability in the colon. No overt signs of disease or inflammation were observed; however, colon length was significantly increased in methanogen-colonized mice compared to controls in both SPF and GF conditions (Extended Data Fig. 1f-g).

**Figure 1.**
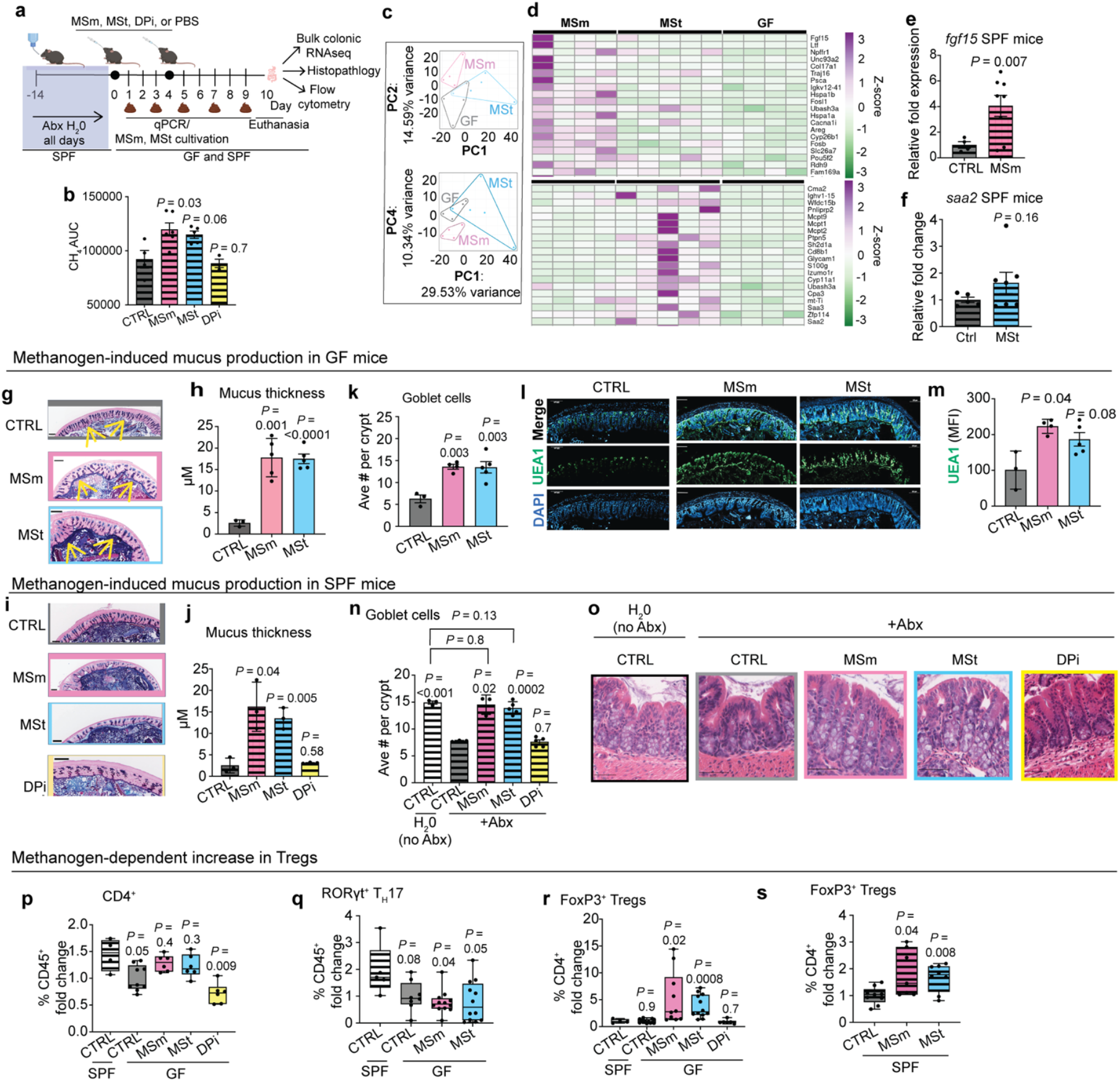
Methanogen colonization promotes epithelial regeneration, mucus production, and regulatory T cell accumulation. **a**, Experimental design. Abx, antibiotics. **b**, Methane concentrations in cultured fecal slurries. AUC, area under the curve. **c**, Principal component analysis (PCA) of colonic transcriptomes from GF, MSm-, and MSt-colonized mice. **d**, Differentially expressed genes (DEGs) of induced transcripts in MSm- or MSt-colonized mice. For (c-d), *n* = 4 (GF, MSm) or 5 mice (MSt). **e–f**, Select transcriptional changes confirmed by RT-qPCR in SPF mice. **g–h**, Representative images (g) and quantification (h) of PAS-AB stained colons from control (CTRL), MSm- and MSt-colonized GF mice. For (g), scale bar = scale bar = 100 µM. **i-j**, Representative images and quantification showing increased colonic mucus thickness in MSm- and MSt-colonized mice compared with SPF or DPi controls. For (i), scale bar = 100 µM. **k–m**, PAS-AB and UEA1 staining demonstrate increased goblet cell abundance in GF mice colonized with MSm or MSt. For (l) scale bar = 100 µM. **n–o**, Goblet cell counts and representative H&E images from SPF mice. For (o), scale bar = 50 µM. **p–s**, Flow cytometric analysis showing the fold change of (p) CD4^+^ cells within the CD45^+^ population, (q) T_H_17 cells within the CD4^+^ population, (r) FoxP3^+^ T_regs_ within the CD4^+^ population in GF mice, and (s) FoxP3^+^ T_regs_ within the CD4^+^ population in SPF mice. For (p-s), box and whisper plots show minimum to maximum values, center denotes median, and the bounds denote the 25th–75th percentiles. For (b, e, f, h, j, k, m, n) data are shown as individual values. Error bars represent the SEM. For (g, i, l, o), at least 25 measurements per mouse were taken. For (b, e, f, h, j, k, m, n, p, q, r, s), *P* values were determined using the Student’s *t*-test with Welch’s correction.

As so little is known about how methanogens shape colonic physiology, we performed RNAseq. Transcriptomic analysis of colonic tissues from GF, MSm- and MSt-colonized mice revealed distinct transcriptional profiles (Fig. 1c): 66 transcripts were increased and 93 were decreased in MSm-colonized mice versus GF; 88 transcripts were upregulated and 38 were downregulated in MSt versus GF (Extended Data Table 1). Of these, 28 transcripts were similarly regulated by MSm and MSt (Extended Data Fig. 2a). In MSm-colonized mice, homeostasis- and proliferation-related transcripts such as *Fgf15* (∼10-fold increase), *Hspa1a, Hspa1b*, and *Fosl1* ^12,13^ were significantly induced (Fig. 1d). Additionally, MSm colonization significantly upregulated *SPDEF* (Extended Data Fig. 2b), which encodes a key regulator of goblet cell differentiation ^14^. In MSt-colonized mice, mast-cell-associated proteases (e.g., *Cma2, Mcpt9, Mcpt1, Mcpt2*) were prominent (Fig. 1d). Gene ontology enrichment indicated up-regulation of processes linked to epithelial integrity (Extended Data Fig. 2c-d) and down-regulation of immune-response pathways (Extended Data Fig. 2e-f) in both methanogen-colonized groups. Some of these transcript changes were recapitulated in SPF mice (Fig 1e-f), although some changes were dependent on GF status (Extended Data Fig. 2g-i). Together, these data suggest that methanogen colonization drives gene programs favoring epithelial maintenance and proliferation.

**Figure 2.**
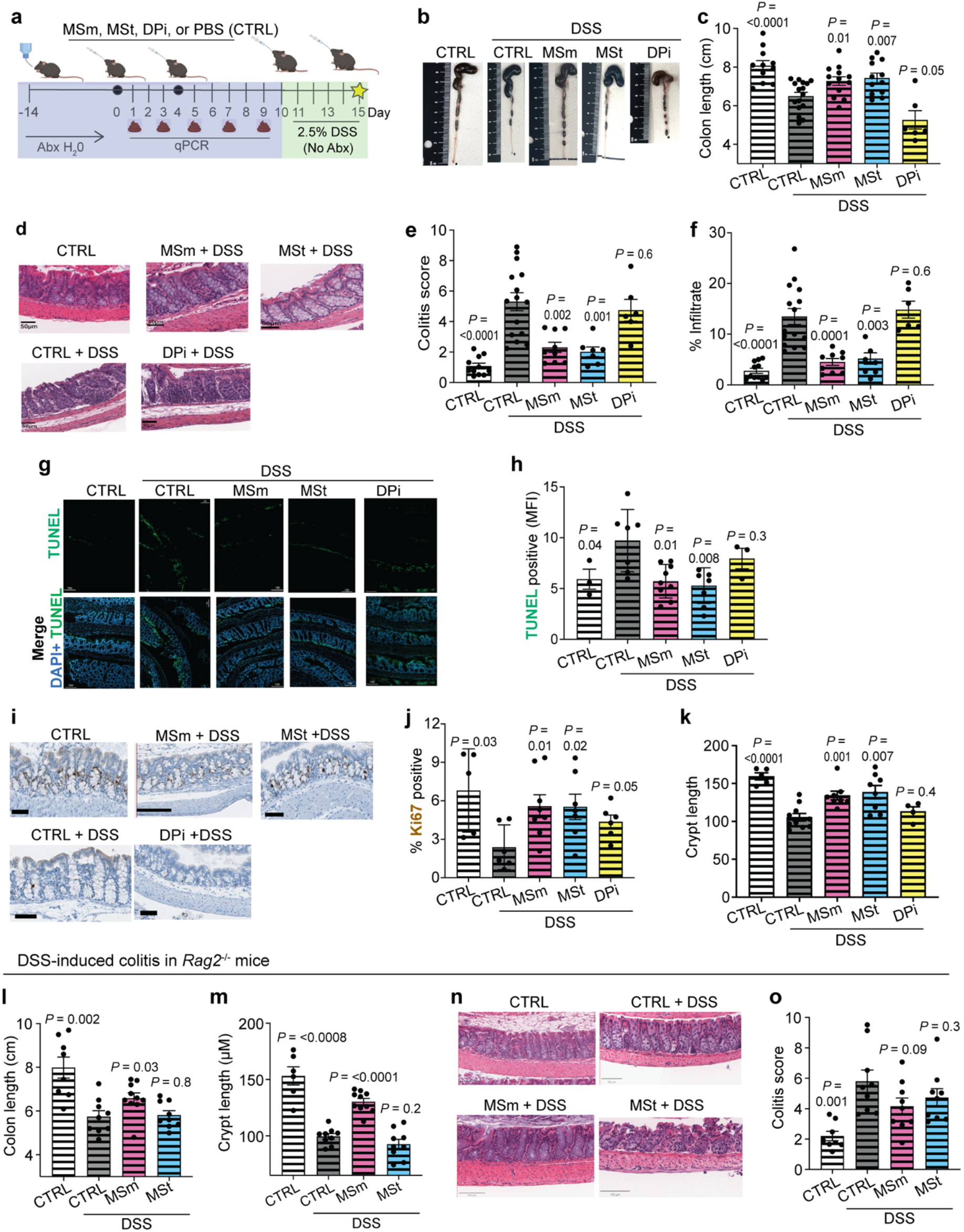
Methanogens protect against DSS-induced colitis through distinct immune mechanisms. **a**, Experimental design: antibiotic-treated SPF mice were colonized for 10 days with *M. smithii* (MSm), MSt, or DPi, then exposed to 2.5% dextran sodium sulfate (DSS). **b**, Representative colon images. **c**, Colon lengths. **d**, Representative H&E staining of colon slices. Scale bars = 50 μM **e**, Histological scoring. **f**, % of inflammatory infiltrate in colon slices quantified using machine learning. **g**, Representative DAPI and TUNEL stained images of colons. Scale bar = 100 µM. **h**, TUNEL positive cells based on mean fluorescence intensity (MFI) averaged from 3-6 fields of view. **i**, Representative images of colonic sections stained to detect Ki67. Scale bar = 50 µM. **j**, Quantification of Ki67 staining. **k**, Crypt lengths based on measurements taken from the base to the apical end of each crypt with ≥ 25 measurements taken per mouse sample. **l**, Colon lengths. **m**, Crypt lengths based on measurements taken from the base to the tip of each crypt with ≥ 25 measurements taken per mouse sample. **n**, Representative H&E staining of colon slices. Scale bars = 50 μM. **o**, Histological scoring. For (b-k), measurements and images were taken at day 5 of DSS-treatment. For (l, m, n, o), measurements and images were taken at day 7 of DSS-treatment. For all, each dot represents one mouse. Error bars show the SEM. *P* values were determined using the Student’s *t*-test with Welch’s correction based on comparison to the CTRL+DSS group.

Mucus is essential to epithelial homeostasis, defense, and repair ^15^ and is shaped by the intestinal microbiota ^16^. MSm- and MSt-colonized mice exhibited a markedly thicker colonic mucus layer compared to GF controls (Fig. 1g-h) or SPF control and DPi-colonized mice (Fig. 1i-j), in which only a thin mucus layer was observed. Quantification of UEA1 lectin confirmed that MSm and MSt increased mucus-producing goblet cells in GF mice (Fig. 1k-m). Goblet-cell counts (H&E staining) confirmed that these findings were recapitulated in SPF mice (Fig. 1n-o). Antibiotic treatment reduced goblet cell numbers and thinned the mucus layer, consistent with impaired secretion (25); these effects were prevented by MSm and MSt colonization but not by DPi (Fig. 1n-o).

Given the links between the microbiota, mucosal integrity, and immune modulation ^17,18^, we analyzed T-cell populations in the colon. Consistent with previous reports ^19,20^, CD4^+^ T-cells were reduced in GF mice, and this reduction was prevented by MSm and MSt, but not by DPi (Fig. 1p). T helper 17 (T_H_17) cell numbers were unaffected by any colonization (Fig. 1q). Notably, Foxp3^+^ regulatory T cells (T_regs_) accumulated significantly in GF and SPF mice that were colonized by MSm and MSt (Fig. 1r-s).

To investigate the biological relevance of methanogen colonization, antibiotic-treated SPF mice were colonized for 10 days with MSm, MSt, or DPi then exposed to 2.5% dextran sodium sulfate (DSS) (Fig. 2a). MSm and MSt colonization prevented significant weight loss and morbidity (Extended Data Fig. 3a-b) and colon shortening (Fig. 2b-c). Moreover, MSm- and MSt-colonized mice exhibited decreased pathology (Fig. 2d-e), suppression of immune cell infiltration (Fig. 2f), and reduced cell death and crypt loss (Fig. 2g-h and Extended Data Fig. 3c). Enhanced epithelial proliferation, measured by Ki67 staining, and crypt lengthening were observed (Fig. 2i-k), consistent with transcriptional and mucus-associated changes. DPi did not confer protection from colitis (Fig. 2b-k, Extended Data Fig. 3a-b).

Because MSm and MSt increased homeostatic T_regs_, we evaluated adaptive immune dependence by repeating the DSS experiments using *Rag2*^− */* −^mice that lack T and B cells. In these animals, MSm colonization still conferred significant protection (morbidity, colon and crypt length) (Fig. 2l-m), though occult blood and moderate histology changes remained (Fig. 2n-o). In contrast, MSt colonization failed to protect *Rag2*^−*/*−^mice on all measured parameters (Fig 2l-o, Extended Data Fig. 3d-e). These data indicate that MSt-mediated protection is lymphocyte-dependent, whereas MSm protection is partly independent of T and B cells.

This study establishes that the gut methanogenic archaea MSm and MSt promote colonic homeostasis by reinforcing the epithelial barrier and limiting inflammatory damage. By colonizing germ-free and bacterial-depleted mice, we show that MSm and MSt promote epithelial regeneration, strengthen the mucus barrier, expand regulatory T cells, and protect against DSS-induced colitis. These findings reveal an unexpected protective role for archaea in gut health. Although MSm and MSt induce distinct transcriptional programs, both enhance epithelial turnover and mucus production, key determinants of barrier integrity. MSm and MSt colonization resulted in T_reg_ accumulation, echoing immune signatures elicited by tolerance-promoting bacteria *Clostridia* sp. and *Bacteroides fragilis* ^18,20^. Yet unlike these bacteria, methanogens lack peptidoglycan and many canonical MAMPs, suggesting that methanogens use unique structural or metabolic cues to engage the mucosal immune system. These may include putative capsule and adhesin-like proteins ^21^. Future studies are necessary to dissect archaeal-derived microbial-associated molecular patterns (MAMPs) and their specific host-cell targets. Understanding these archaeal signals will be critical for defining the mechanisms of host–archaea communication.

Our functional analyses further show that MSm and MSt protect from colitis through nonredundant pathways. MSm retains partial efficacy in lymphocyte-deficient mice, indicating contributions from epithelial or innate responses, whereas MSt requires adaptive immunity. These findings also reconcile earlier tissue culture ^22,23^ or murine airway inflammation models ^24,25^ that reported pro-inflammatory effects of non-viable methanogens or methanogen cellular components, by demonstrating that live archaea in the native intestinal niche elicit protective rather than inflammatory responses.

Translationally, methanogens or their derived factors may offer a strategy to strengthen the epithelial barrier and curb chronic inflammation. Given that chronic inflammatory disorders such as IBD increase colorectal cancer risk, methanogen colonization might also limit tumorigenesis. Albeit the observed stimulation of epithelial proliferation suggests a potential caveat that warrants further investigation. Together, our findings establish human-associated methanogenic archaea as active modulators of epithelial and immune function and underscore the need to identify their molecular effectors and community-level interactions to define how these ancient symbionts shape intestinal and systemic health.

## Supporting information

RNAseq data

## Acknowledgments

This research was supported by NIH grants to M.M.K. (AI162696 and AI146888). An NIH Biodefense & Infectious Diseases Short-Term Training to Increase Diversity in Biomedical Research (T35AI060528) provided support for B.W.H. This work was further supported by the University of Virginia Trans-University Microbiome Initiative (TUMI) Award to M.M.K. and a UVA MIC Innovation Award to M.M.K. and M.R.R. The Research Histology Core (RRID): SCR_025470, the Biorepository and Tissue Research Facility (RRID): SCR_022971, and the Flow Cytometry Core are supported by UVA School of Medicine (RRid:SCR_017829). The Flow Cytometry Core received additional support through the NCI (P30-CA044579). We thank H. Agaisse, Z. Lifschin, S. Rolland, N. Jadeja, J. Kashatus, K. Thomas, E. Lamb, and M. Wakim Broden for sharing protocols and helping with troubleshooting and technical aspects of the microscopy experiments. We thank the UVA Germ-Free Facility for assistance with the gnotobiotic mouse experiments.

## Contributions

E.E.B designed, conducted, and analyzed experiments and wrote the paper; W.E.W., S.H.K., A.M.P., and M.T.M. conducted experiments; B.W.H, M.D.S., and M.M.R. designed and analyzed experiments; H.Z. analyzed experiments; M.M.K. supervised all experiments, analyzed data, and wrote the paper.

