## Supplementary material for "Methanogenic archaea bolster mucosal homeostasis and protect from colitis": Methods

**Microbial strains and culture conditions.** The anaerobic culture techniques of Hungate [1], as modified by Sowers and Noll [2], were used in this study. *Methanobrevibacter smithii* (MSm) type strain PS and *Methanosphaera stadtmanae* (MSt) strain MCB-3 were purchased from the DSMZ-German Collection of Microorganisms and Cell Cultures GmbH and were routinely grown in modified methanobacterium medium 1523 (dsmz.de) with the addition of 10% rumen fluid (Bar Diamond), with an overpressure of 1 atm $H_2$ [3]. MSt culture medium was also supplemented with 5% methanol (Fisher, MMX04904). DPi strain VPI C3-23 was purchased from the American Type Culture Collection (ATCC 29098) and was grown in Tryptic Soy Broth Fisher, DF0370-17-3) supplemented with 0.5 g/L $Na_2S$ (Fisher, AA6512222) and 5% defibrinated sheep's blood (Fisher, R54016). *Bacteroides thetaiotaomicron* strain VPI-5482 was purchased from the ATCC (ATCC 29148D-5) and was grown in modified methanobacterium medium 1523, supplemented with 10 mM glucose (Acros Organics, 492-62-6). All cultures were grown at 37$^o$C with shaking.

**Methane analysis.** Methane from microbial cultures and cultured fecal pellets was measured using a Shimadzu gas chromatograph 2014. For this, ~200 $\mu$L of headspace was sampled from culture vials. Methane levels were determined by calculating area under the curve during PostRun analysis using LabSolutions software.

**RNA and DNA purification.** Nucleic acids were extracted using the PureLink RNA Mini Kit (Invitrogen). Samples for RNAseq and RT-qPCR were treated with DNase I (Millipore Sigma). Samples for qPCR were treated with RNase I (Millipore Sigma). All samples were ethanol precipitated prior to use. RNA purity was determined by measuring the $A_{260}/A_{280}$ absorbance ratio and by performing PCR (35 cycles).

**RT-qPCR.** RT-qPCR was performed in a one-step reaction using an ABI 7500-FAST sequence detection system and software (Applied Biosystems) [4]. For each 10-µl reaction mixture, 5 µl 2× SYBR master mix (Invitrogen, 4367659), 0.05 µl Multi-Scribe reverse transcriptase (Invitrogen, 4308228), and 0.05 µl RNase inhibitor (Invitrogen, N8080119) were added. Primers were designed using Primer Blast (NCBI) to ensure no cross-reactivity and are listed in Supplementary Information Table 1. Amplicon length was approximately 100 bp. Amplification efficiency of each primer pair was verified using standard curves of known DNA concentrations. Melting-curve analysis was used to ensure template specificity by heating products to 95ºC for 15 s, followed by cooling to 60°C and heating to 95ºC while monitoring fluorescence. After the amplification efficiency and template specificity were determined for each primer pair, relative quantification analysis was used to analyze the samples using the following conditions for cDNA generation and amplification: 1 cycle at 48ºC for 30 min, 1 cycle at 95ºC for 10 min, and 40 cycles at 95ºC for 15 s and 60ºC for 1 min. Two technical replicates of each biological replicate were included for each gene target. Data were normalized to the reference controls *Rplp0* or *eef2,* and analyzed using the comparative critical threshold ($C_T$) method [5]. The expression level of the target genes was compared using the relative quantification method [5]. Data are presented as the change (*n*-fold) in expression levels compared to control (CTRL) levels. Error bars represent the standard deviations of the $\Delta\Delta C_T$ value.

**Microbial enumeration for mouse engraftments.** An autofluorescence and SYBR Gold flow-cytometry based method based on [6] was used to quantify MSm, MSt, and DPi cell number/ mL prior to mouse engraftment. A 1000 $\mu$L aliquot of exponentially growing MSm, MSt, or DPi culture was fixed in 4% formaldehyde (Fisher, BP531-500) and incubated for 10 min at room temperature. Then, samples were re-pelleted and suspended in 200 $\mu$L of SYBR Gold reagent (Thermo Fisher, S11494) for 45 min at 4°C. Cells were washed and then resuspended in phosphate-buffered saline (PBS) (Gibco, 10010023) for flow cytometry. Samples were analyzed using an Aurora Northern Lights cytometer. Controls included fixed, unstained MSt ($F_{420}$ (autofluorescence) reference); unstained *B. thetaiotaomicron* (negative control), and SYBR Gold-stained *B. thetaiotaomicron* (single-positive control). Cells were gated on FSC/SSC, $F_{420}^+$, $F_{420}^+$/SYBR Gold$^+$. SpectroFlo software was used for compensation with live unmixing and autofluorescence subtraction, double-positive cells (methanogens) were quantified using FlowJo software version 10.10.0.

**Animal studies**. All animal work was approved by the University of Virginia Institutional Animal Care and Usage Committee (IACUC). Germ-free C57BL/6 aged 4-5 weeks (Taconic) were housed in sterile isolators at the germ-free facility at the University of Virginia School of Medicine. Upon arrival and throughout the experiments, mice were routinely tested by qPCR and culturing to confirm germ-free/ mono-association status. Conventional C57BL/6 (Jax 000664) and *Rag2*$^{-/-}$ (Jax 008449) aged 3 weeks were purchased from the Jackson Laboratory. Conventional mice were given filter-sterilized water containing ampicillin (1 g/L; Fisher, BP1760), vancomycin (0.5 g/L; Thermo Fisher, J6279006), gentamicin (0.05 g/L; Millipore Sigma, 48760), streptomycin (1 g/L; Thermo Fisher, 45341000), tetracycline (0.5 g/L; Fisher, BP912-100), and sucrose (4 g/L; Millipore Sigma, S0389) for 14 days prior to and throughout microbial colonization. For homeostasis experiments, mice were euthanized on day 10 post engraftment. For the DSS-colitis experiments, antibiotics were removed from the drinking water immediately prior to DSS administration as antibiotics can influence the severity DSS-induced colitis [7,8].

**Microbial intestinal colonization and enumeration.** Mice were orally gavaged with $1 \times 10^8$ microbial cells on days 0 and 4. To confirm colonization, fecal pellets were collected from control, MSm-, MSt-, and DPi-colonized mice at 1, 3, 5, 7, and 9 days after the initial gavage. DNA was extracted as described above. qPCR was performed using MSm-, MSt-, and DPi-specific 16S rRNA gene primers (listed in Supplementary Information Table 1). rRNA gene copies/ g of feces was quantified as described [9].

**Cell isolation and flow cytometry.** Colons were gently harvested by cutting at the cecum-colon junction and immediately processing the tissue. Unless indicated, fecal content was removed, and colons were flushed with PBS. Tissues were flushed with PBS and fat was removed. Subsequently, tissues were treated with HBSS (Gibco, 14175095) with 5 mM EDTA (Boston Bioproducts, BM-150), 1 mM DTT (Thermo Fisher, R0861), 5% FBS (Optima, 512450), and 20 mM HEPES (Gibco, 15630-080) for 20 min at 37°C with gentle agitation and then vortexed (15-20 s). This was performed twice. Then, tissues were washed in PBS, minced, and digested in a 5% FBS solution containing collagenase (0.5 mg/ml; Millipore Sigma, 11088858001) and DNase I (0.5 mg/ml; Millipore Sigma Millipore Sigma, 10104159001) and 20 mM HEPES. To make single-cell suspensions, the digested tissues were passed through 100 µm cell strainers (Corning, 352360) using mechanical force with the rubber end of a 5 mL syringe.

Cells were stained with the Zombie Aqua Fixable Viable Kit (Biolegend, 423101) followed by staining for surface markers. Cells were then fixed with 1% methanol formaldehyde solution, followed by permeabilization in 0.5% Saponin solution (both from Thermo Scientific, 00-5523-00) and intracellular staining. SPHERO™ AccuCount Particles (Fisher Scientific, NC9643447) were utilized to enumerate cell counts. Flow cytometry experiments were performed on a Cytek Aurora Borealis (5 lasers). Antibodies and vendors are listed in Supplemental Information Table S2. Flow gating strategy is shown in Supplemental Information Figure 1. Flow Cytometry data was analyzed using FlowJo version 10.10.0.

**Histology.** For assessment of colonic mucus and goblet cells, intact colons were fixed overnight at 4°C in Carnoy's fixative (dry methanol:chloroform:glacial acetic acid in the ratio 60:30:10) (Fisher, C298-4 and BP1185-500) and subsequently transferred to 70% ethanol (Fisher, BP2818500) [10]. For all other histopathological analyses, colonic tissues were cut open longitudinally and displayed in cassettes as swiss-rolls. Cassettes were immersed in 10% neutral buffered formalin [11]. For paraffin slides, the tissue was preserved in 70% ethanol before loading onto a tissue processor for dehydration and paraffin infiltration by the UVA Research Histology Core. After manual embedding into a paraffin block, paraffin sections were cut at 5 μm on a Leica microtome.

**Histopathology analyses.** To determine the histopathology colitis score (HCS), H&E-stained samples were de-identified prior to analysis by a blinded reviewer. Samples were assessed from the muscularis mucosae using an annotation feature in Qupath [12], where sections of intestinal mucosa were designated either normal, injured, or ulcered [13] (see below). The HCS was calculated as the sum of percent injured and two times the percent ulcered divided by ten [13].

| Colitis scoring criteria | Representative Image |
|---|---|
| Normal<br>• No leukocyte infiltrate<br>• Crypts arranged parallel<br>• Goblet Cells intact<br>• Colonocytes intact<br>• Muscularis mucosae intact | 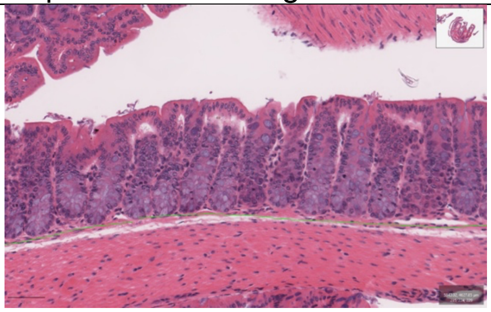<br>Scale Bar = 50 μm |
| Injured<br>• Inflammatory Infiltrate<br>• Crypt Dysplasia<br>• Crypt hyperplasia<br>• Goblet cell loss | 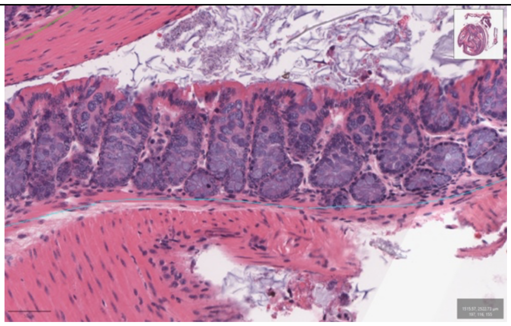<br>Scale Bar = 50 μm |
| Ulcered<br>- Absence of Crypt structure<br>- Leukocyte Infiltrate<br>- Ulcerated from muscularis mucosae to brush border | 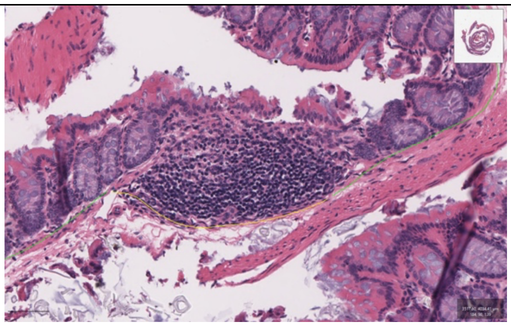<br>Scale Bar = 50 μm |

**Infiltrate quantification:** To obtain infiltrate quantification from colitis samples, imaged swiss rolls were assessed using the cell detection feature in Qupath to produce an undifferentiated cell count. To enumerate the percent infiltrate, tissue was manually annotated based on cell size and color intensity (see below). A total of 20-50 measurements were taken within each sample. The percent infiltrate in every sample was calculated as number of infiltrating cells divided by total cells.

| Cell Counter Designation | Representative Example |
|---|---|
| Leukocyte Infiltrate | 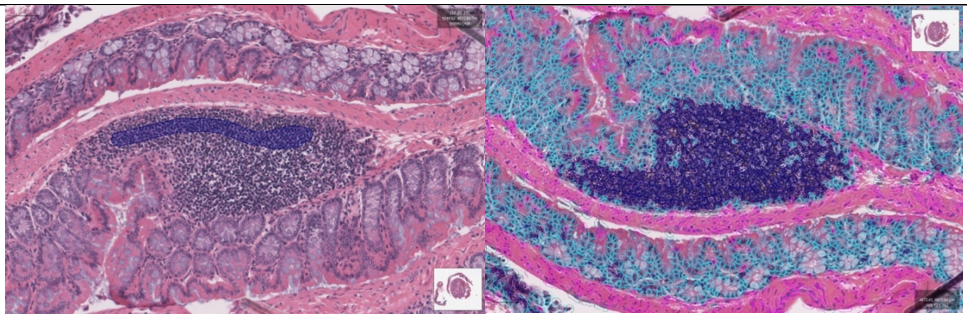 Scale Bar = 50 μm |

**Immunofluorescence microscopy.** Sample preparation was performed based on methods described in [11]. Paraffin sections were re-hydrated in the following sequence: xylene (Fisher, X3P-1GAL**)**, 100% ethanol, 95% ethanol, 70% ethanol, and water. Antigen retrieval was performed at 95–100 °C for 20 min using a pre-heated citric acid-based buffer (Vector Laboratories, H-3300). Slides were cooled in the buffer solution for 1 h at room temperature and washed in PBS 2Å~ for 5 min each. Slides were permeabilized in 0.1% Triton for 10 min at room temperature and washed in PBS for 5 min. Slides were cooled in the buffer solution for 1 h at room temperature and washed in PBS 2% for 5 min each. Slides were permeabilized in 0.1% Triton (Fisher, 648466) for 10 min at room temperature and washed in PBS for 5 min. Samples were blocked in 5% bovine serum albumin (Millipore Sigma, A9647-50G) and 2% normal goat serum (Thermo Fisher, 50197Z) in PBS for 1 h at room temperature. Primary antibodies were diluted 1:100 in blocking buffer (E-cadherin, Thermo Fisher 20874-1-AP) (Ulex Europaeus (Gorse) Agglutinin I (UEA I), L32476, Thermo Fisher) and incubated overnight at 4 °C. Secondary antibodies (goat anti-rabbit Alexa Fluor™ 647, A-21245 Thermo Fisher Scientific) were diluted 1:500 in blocking buffer and incubated for 2h at room temperature.

All immunofluorescence experiments were imaged using a Zeiss LSM 900 confocal microscope equipped with Airyscan. ZEN Blue 3.3 software was used for image acquisition and processing, including scale bar generation. Image J Fiji was utilized for immunofluorescent image quantification.

**TUNEL assays.** The TUNEL assay was performed according to the manufacturer's instructions (In situ death detection kit, TMR red, Roche, 12156792910). Slides from immunofluorescence experiments were stained with DAPI ((4′,6-diamidino-2-phenylindole, D1306, Thermo Fisher), according to manufacturer instructions. Coverslips were mounted using ProLong Gold Antifade Mountant (Thermo Fisher, P36934). A total of 3–6 representative images per tissue sample were analyzed and averaged to determine mean fluorescence intensity (MFI) of the AF488 channel. The average TUNEL+ MFI quantification for each mouse sample was reported.

**UEA1 quantification.** UEA1 was measured to determine goblet cell numbers per sample. A total of 14-15 randomly selected UEA1-FITC stained crypts per tile image were analyzed (3–5 tile images per mouse). The average goblet cell number per mouse colon was calculated. To quantify UEA1+ MFI, 40-50 goblet cells per randomized tile image were counted (3-5 tile images per mouse). The average UEA1+ MFI of surface area per mouse colon was reported.

**Mucus imaging and quantification.** Post fixation, tissues were paraffin-embedded (described above) and stained with Alcian Blue/Periodic Acid Schiff (AB/PAS) or hematoxylin and eosin (H&E). Slides were imaged using a Aperio ImageScope Scanscope scanning microscope and analyzed using QuPath software. AB/PAS-stained slides were imaged for mucus quantification. For this, 25 randomized mucus thickness quantifications were measured per mouse colon (3-5 colon cross sections per colon). H&E-stained slides were imaged for goblet cell enumeration. For 15 randomized sections were assessed. For mucus and goblet cells, the average per mouse sample was plotted.

**Brightfield microscopy and quantification.** Tissue samples were processed were stained with Periodic Acid Schiff/Alcian Blue (PAS/AB) or hematoxylin and eosin (H&E) by the UVA Research Histology Core. Brightfield experiments were imaged using a Aperio ImageScope Scanscope scanning microscope and analyzed using QuPath software.

**Mucus and goblet cells.** For PAS/AB-stained slides, 25 randomized mucus thickness quantifications were measured per mouse colon (3-5 colon cross sections per colon). For H&E-stained slides 15-25 randomized crypts were quantified for the # of goblet cells/crypt. For mucus and goblet cells, the average per mouse sample was plotted.

**Crypt length.** H&E-stained samples were quantified by measuring approximately 50 randomized measurements per mouse colon swiss-roll. The average crypt length per mouse sample was plotted.

**RNAseq and analysis.** RNA purified from whole colons was barcoded and sequenced by the University of Maryland Genomic Research Core. After rRNA depletion, sequencing libraries were generated and sequenced on an Illumina Novaseq platform. Reads were trimmed and mapped to the *Mus musculus* genome and HTSeq [14] was used for read counting and FPKM (Fragments Per Kilobase of transcript sequence per Millions base pairs sequenced) calculation. Pseudogenes were excluded from subsequent analyses based on Ensembl gene annotation (GRCm39, release 111). For principal components analysis (PCA), raw read counts were normalized using the median ratio method implemented by DESeq2 [15,16], PCA was then performed in R using the prcomp() function. The differential expression analysis was performed using DESeq2. Genes with absolute log2 fold change > 0.8, *P* value < 0.05, and FDR < 0.05 compared to GF mice were considered differentially expressed. Gene ontology-based functional enrichment analysis was performed using David NCBI tool [17]. Ensembl Mouse IDs were converted to Entrez IDs using the MGI mouse database. All the related plots were generated using ggplot2 [18].

**Statistical Analyses**

Unless indicated, differences between the experimental groups were analyzed using the student's *t* test. Data was analyzed using GraphPad Prism.

**Supplemental Information Table 1. Primers used in this study.**

| Purpose | Primer Name | Sequence (5' to 3') |
|---|---|---|
| PCR | F: MS_RT_UPF1: | AACGAGTTTAGGCTCGTGGC |
| PCR | R: MS_RT_DOWNR1 | CAATCCCTTTTTGGGTGGCG |
| PCR | F: MStadt_16Sup: | ATCTGCGGCTGATTAGGTCG |
| PCR | R: MStadt_16Sdown: | CCAGGAGATTCGGGGCATAC |
| PCR | F: 93_dp16S_bigF1: | ACCAAGGCAACGATGGGTAG |
| PCR | R: 94_dp16S_bigR1 | CCCAACATCTCACGACACGA |
| PCR | F: musGAPDH_F1: | TGTAGTGAGCCCCAGGCTAT |
| PCR | R: musGAPDH_R1: | TCCGCCCTGATCTGAGGTTA |
| qPCR | F: M. smit_16S-740F | CCGGGTATCTAATCCGGTTC |
| qPCR | R: M. smit_16S-862R | CTCCCAGGGTAGAGGTGAAA |
| qPCR | F: M.stadt_16S_1F | ATCTGCGGCTGATTAGGTCG |
| qPCR | R: M.stadt_16S_1R | CTCTTGCTCTCACAACCCGT |
| qPCR | F: 91_dp16SqpcrF1 | AGGAACATCAGTGGCGAAGG |
| qPCR | R: 92_dp16SqpcrR1 | GTTTACGGCGTGGACTACCA |
| RT-qPCR | Mus_Eef2_F1 | GCCATGTGTGGTGAAAGAGG |
| RT-qPCR | Mus_Eef2_R1 | TTGTCACAGGACATTGTTGCT |
| RT-qPCR | Mus_ Rplp0 _F1 | ACTGGTCTAGGACCCGAGAAG |
| RT-qPCR | Mus_ Rplp0 _R1 | TCAATGGTGCCTCTGGAGATT |
| RT-qPCR | Mus_18S_F1 | AGTCCCTGCCCTTTGTACACA |
| RT-qPCR | Mus_18S_R1 | CGATCCGAGGGCCTCACTA |
| RT-qPCR | muF7_F1 | AGATGGATAGTGACCGCAGC |
| RT-qPCR | muF7_R1 | GGGCCATACCTACCCATCA |
| RT-qPCR | mus_FGF15_F1 | CGGTCGCTCTGAAGACGATT |
| RT-qPCR | mus_FGF15_R1 | GAACACTCACCAGCCCGTAT |
| RT-qPCR | muSaa1_F1 | TGGTGAGTAGCTTCATCCTGC |
| RT-qPCR | muSaa1_R1 | CAGGAGGCACTGCTAGGACA |
| RT-qPCR | muSaa2_F1 | TATGATGCTGCCCAAAGGGG |
| RT-qPCR | muSaa2_R1 | AGAAGAGTCGAGCCCTGCTA |
| RT-qPCR | muIghv1-74_F1 | GTAGGCTGTGCTGGAGGATTT |
| RT-qPCR | muIghv1-74_R1 | AGGCCTTGAGTGGATTGGAA |
| RT-qPCR | mus_Ighv 1-4_F | AGTTGCATGTAGGCTGTGCT |
| RT-qPCR | mus_Ighv 1-4_R | TGGACAGGGTCTGGAATGGA |

**Supplemental Information Table 2. Antibodies for flow cytometry**

| Marker | Fluorochrome | Clone (as applicable) | Reactivity | Catalog # | Vendor |
|---|---|---|---|---|---|
| FC Block (CD16/32) | - | 93 | Mouse | 101301 | Biolegend |
| CD45R | BUV661 | 30-F11 | Mouse | 612975 | BD Biosciences |
| CD3e | BV785 | 17A2 | Mouse | 100231 | Biolegend |
| CD4 | AF700 | GK1/5 | Mouse | 100429 | Biolegend |
| CD8b | BV480 | H35-17.2 | Mouse | 746835 | BD Biosciences |
| Foxp3 | Pac Blue | MF-14 | Mouse | 126409 | Biolegend |
| Ror$\gamma$T | PE | AFKJS-9 | Mouse | 12-6988-82 | Invitrogen/Thermo Fisher |

**Supplemental Information Figure 1.** Flow cytometry gating strategy.

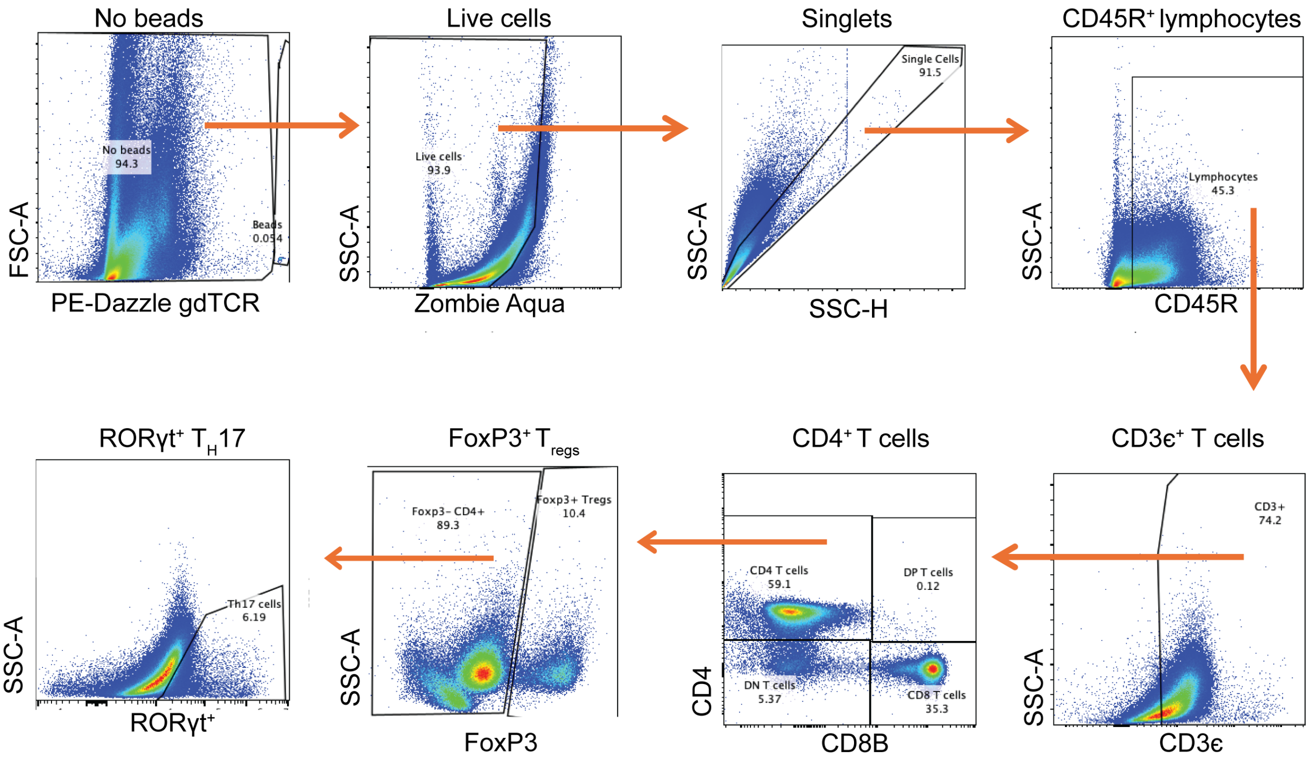
