## Extended Data Figures for "Methanogenic archaea bolster mucosal homeostasis and protect from colitis"

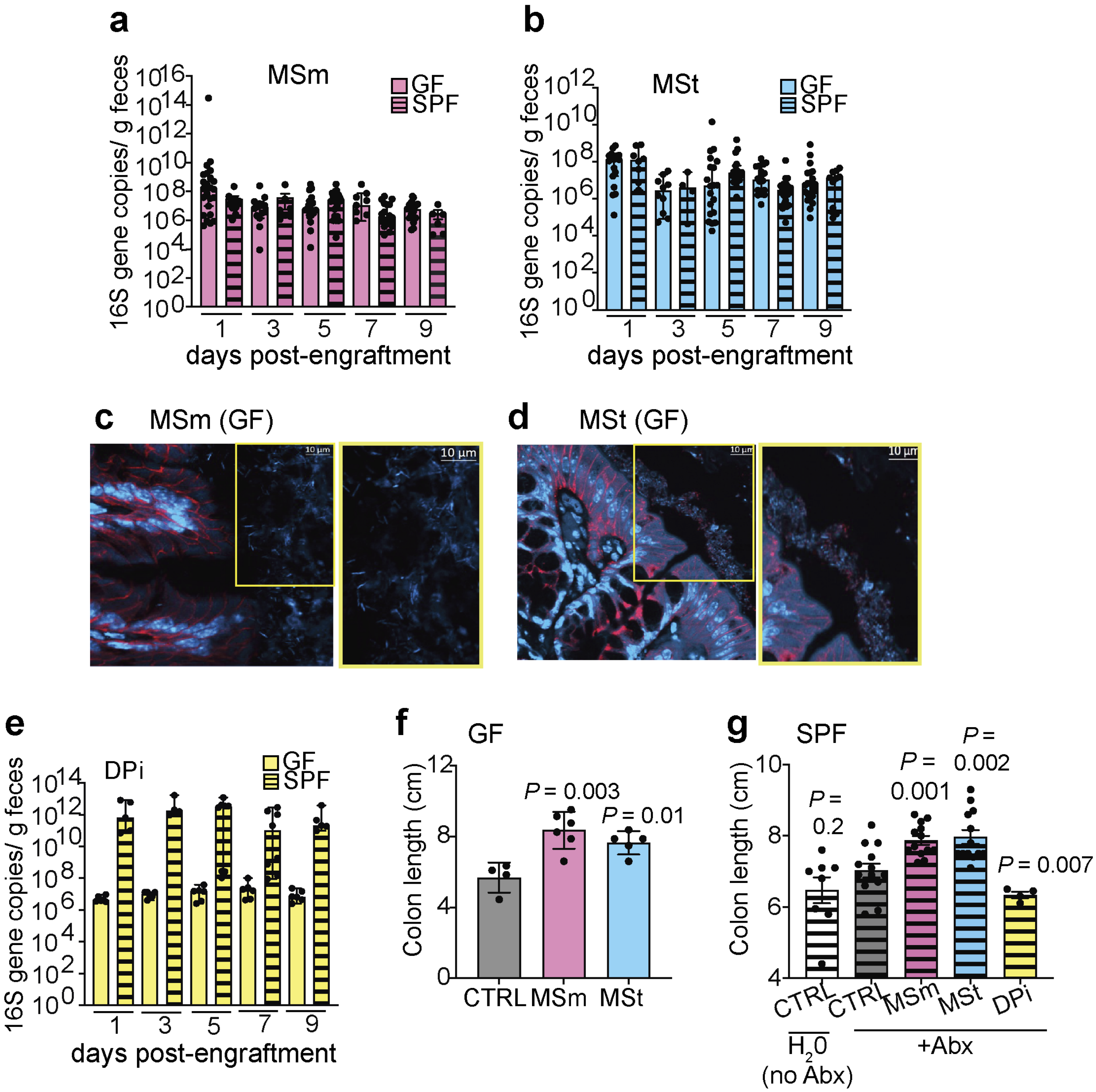


**Extended Data Figure 1. Methanogen colonization of GF and SPF mice. a-b**, Fecal qPCR confirming stable colonization of MSm and MSt in GF (solid bars) and SPF (striped bars) mice. **c–d**, Representative images of colonic sections showing localization of MSm and MSt to the mucus layer in GF mice. Scale bars = 10 µM. **e**) Fecal qPCR confirming stable colonization of the niche-matched control DPi in GF (solid bars) and SPF (striped bars) mice at. **f–g**, Colon lengths from (f) GF and (g) SPF mice at 10 days post-engraftment. For (a, b, e), data are shown as individual values, error bars represent and median with 95% CI. For (f, g), error bars represent the SEM. *P* values were determined using the Student's *t*-test with Welch's correction.

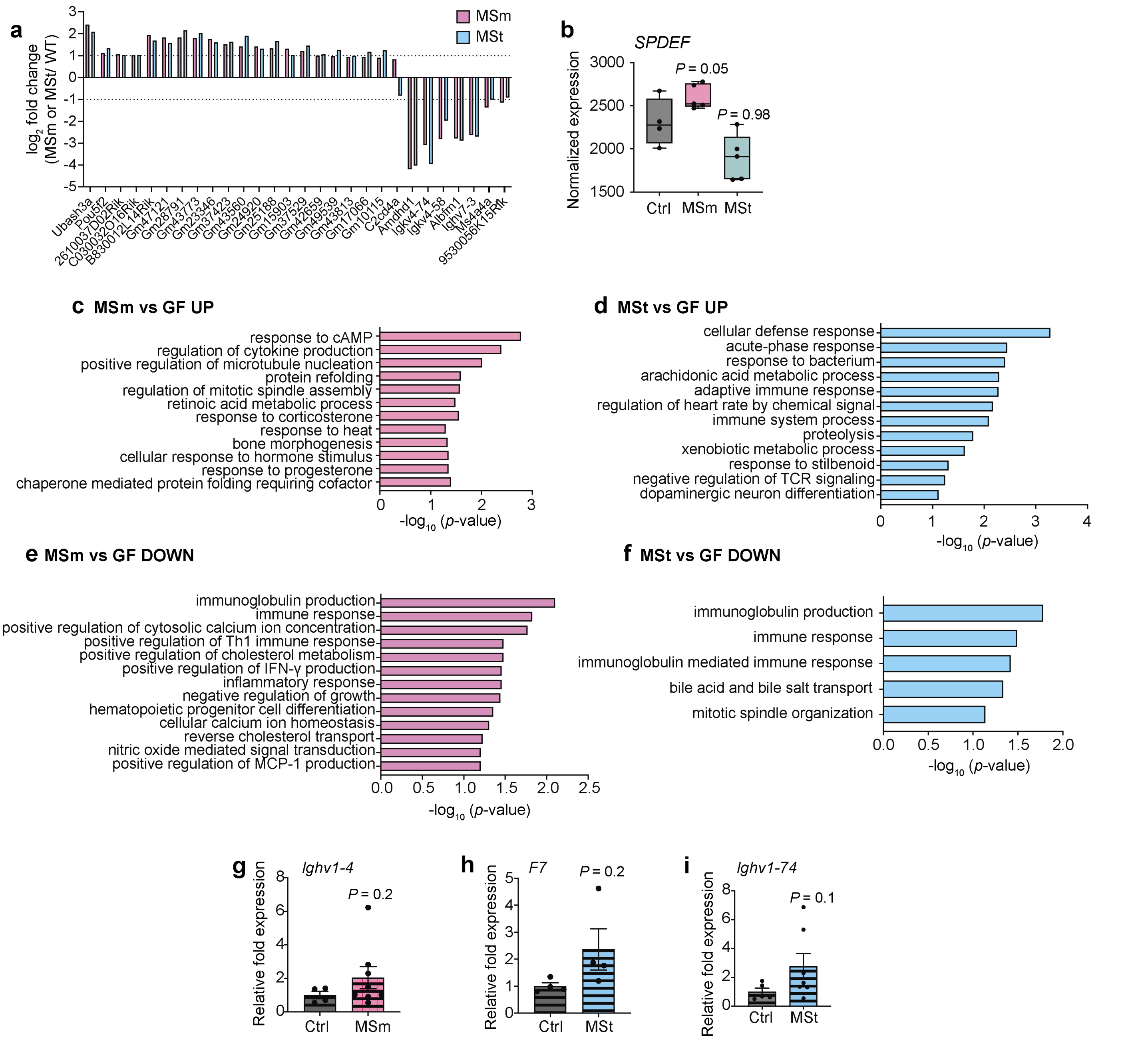


**Extended Data Figure 2. Transcriptomic changes in colonic tissues following methanogen colonization.** **a**, RNAseq data of differentially expressed genes (DEGs) that were common in MSm- and MSt-colonized GF mice relative to GF controls. **b**, RNAseq data showing up-regulation of *SPDEF*, a key transcription factor for goblet cell differentiation, in MSt-colonized mice. *P*-values determined by Wilcoxon Rank Sum test. **c–d**, Gene ontology (GO) enrichment analysis highlighting up-regulation of processes associated with epithelial integrity and proliferation in (c) MSm-colonized mice and (d) MSt-colonized mice. **e–f**, GO enrichment analysis highlighting down-regulation of immune-response pathways in (e) MSm- and (f) MSt-colonized mice. **g–i**, Select transcript changes confirmed in SPF mice. For data shown in (a, b, c, d, e, f), $n$ = 4 (GF, MSm) or 5 mice (MSt). For (g-i), data are shown as individual values, error bars represent the SEM, and *P* values were determined using the Student's *t*-test with Welch's correction.

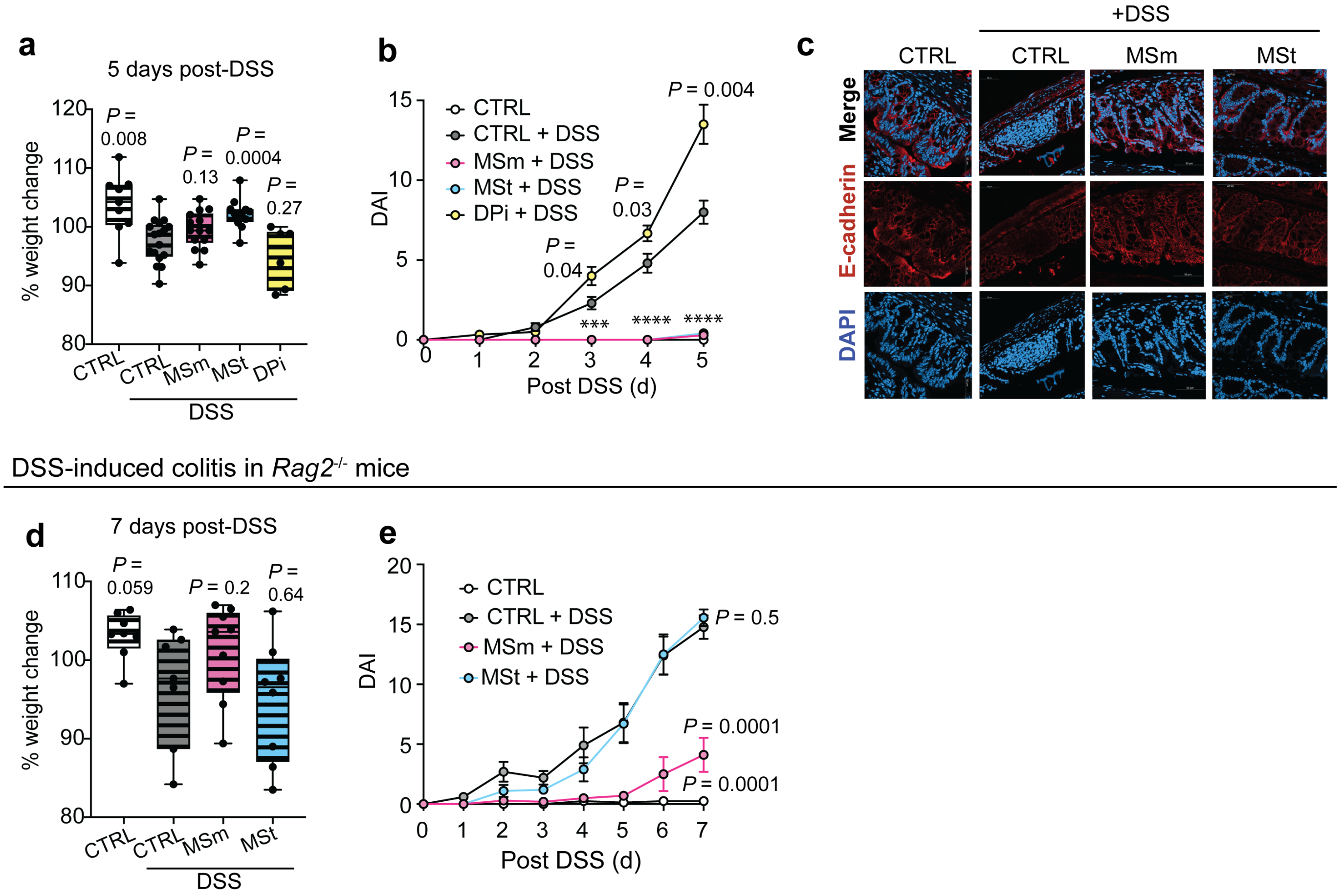


**Extended Data Figure 3. Methanogens protect from DSS-induced inflammation and epithelial damage. a,** Body weight and **b,** disease activity index (DAI) of SPF mice colonized with MSm, MSt, or DPi and challenged with 2.5% DSS. **c**, Representative images of colonic sections showing increased epithelial integrity and decreased crypt loss and ulceration in MSm- and MSt-colonized DSS-treated mice compared to control (CTRL) and DPi-colonized DSS-treated mice. Scale bars = 50 μM. **d,** Body weight and **e,** disease activity index (DAI) of antibiotic-treated *Rag2*$^{-/-}$ mice. Data are shown as individual values. Error bars represent the SEM. *P* values were determined using the Student's *t*-test with Welch's correction. *** *P* = 0.0003, **** *P* = 0.0001.
